# Epilepsy-associated Mutations in the Calcium-sensing Receptor Disrupt the Regulation of NALCN Sodium-leak Channel by Extracellular Calcium in Neurons

**DOI:** 10.1101/2020.11.07.372623

**Authors:** Chunlei Cang, Boxun Lu, Dejian Ren

**Affiliations:** Department of Biology, University of Pennsylvania, Philadelphia, PA 19104, U.S.A; University of Science and Technology of China, Hefei, Anhui 230026, China; State Key Laboratory of Medical Neurobiology and MOE Frontiers Center for Brain Science, School of Life Sciences, Fudan University, Shanghai 200433, China

## Abstract

Most mammalian neurons have a resting membrane potential (RMP) of ~ −50 mV to −70 mV, significantly above the equilibrium potential of K^+^ (E_K_) of ~ −90 mV. The resting Na^+^-leak conductance is a major mechanism by which neurons maintain their RMPs above E_K_. In the hippocampal neurons, the TTX-insensitive, voltage-independent Na^+^ leak is mediated by the NALCN cation channel. Extracellular Ca^2+^ (Ca^2+^_e_) also controls the sizes of NALCN current (I_NALCN_) in a G-protein-dependent fashion. The molecular identities of the basal Na^+^ conductances and their regulation in other regions in the central nervous system and in the peripheral nervous system are less established. Here we show that neurons cultured from mouse cortices, ventral tegmental area, spinal cord and dorsal root ganglia all have a NALCN-dependent basal Na^+^-leak conductance that is absent in NALCN knockout mice. Like in hippocampal neurons, a decrease in [Ca^2+^]_e_ increases I_NALCN_. Using shRNA knockdown, we show that the regulation of I_NALCN_ by Ca^2+^_e_ in neurons requires the Ca^2+^-sensing G-protein-coupled receptor CaSR. Surprisingly, the functional coupling from [Ca^2+^]_e_ to NALCN requires CaSR’s distal C-terminal domain that is dispensable for the receptor’s ability to couple [Ca^2+^]_e_ to its canonical signaling targets such as PLC and MAPK. In addition, several epilepsy-associated human CaSR mutations, though sparing the receptor’s ability to sense Ca^2+^_e_ to maintain systemic [Ca^2+^], disrupt the ability of CaSR to regulate NALCN. These findings uncover a unique mechanism by which CaSR regulates neuronal excitability via NALCN in the central and peripheral nervous system.

## Introduction

Calcium regulates cellular processes ranging from muscle contraction, cell migration, hormone secretion, neurotransmitter release to gene expression ((Berridge et al., 2003; Clapham, 2007) for reviews). The cytosolic [Ca^2+^] ([Ca^2+^]_cyt_) is tightly regulated, with the resting concentration to be <100 nM. Increases in [Ca^2+^]_cyt_ can be achieved by Ca^2+^ releases from intracellular Ca^2+^ stores such as endoplasmic reticulum and mitochondria, and/or by Ca^2+^ influx from the extracellular milieu. How intracellular Ca^2+^ serves as a “second messenger” to regulate the wide spectrum of physiological process has been extensively studied ((Berridge et al., 2003; Clapham, 2007) for reviews).

In contrast to that of intracellular Ca^2+^, how extracellular Ca^2+^ (Ca^2+^_e_) signals to control cellular function is much less understood ((Brown et al., 1993; Hofer and Brown, 2003) for reviews). The systemic extracellular Ca^2+^ concentration ([Ca^2+^]_e_) is generally regulated to be ~ 1.2 mM in mammals. However, the concentration in the brain can fluctuate significantly, particularly during extensive neuronal activities in regions where extracellular space is limited, and [Ca^2+^]_e_ can drop to as low as 0.1 mM (Benninger et al., 1980; Heinemann and Pumain, 1980; Krnjevic et al., 1980; Nicholson et al., 1977; Pumain and Heinemann, 1985). Changes of >0.2 mM in cerebral cortex [Ca^2+^]_e_ have been observed between sleep and awake in cats and mice (Amzica et al., 2002; Ding et al., 2016). Large variations are also found under pathophysiological conditions such as hypocalcemia, hypoxia, ischemia, trauma and seizure (Brown and MacLeod, 2001; Heinemann et al., 1986; Morris and Trippenbach, 1993; Nilsson et al., 1993; Silver and Erecinska, 1990).

A key player in the maintenance of systemic Ca^2+^ levels is CaSR, a Ca^2+^-sensing G-protein coupled receptor ((Brown et al., 1993; Hofer and Brown, 2003) for reviews). In the parathyroid glands, CaSR detects the systemic [Ca^2+^] level and couples it to the secretion of parathyroid hormones (PTHs), which in turn regulate the balance among Ca^2+^ absorption/reabsorption through the gastrointestinal tract and the kidney, and Ca^2+^ absorption and releases in the bones ((Hebert and Brown, 1996) for a review). Both gain-of-function and loss-of-function CaSR mutations have been discovered and are implicated in the malregulation of systemic Ca^2+^ levels (Dershem et al., 2020; Pidasheva et al., 2004).

Intriguingly, CaSR is also widely expressed throughout the brain (Allen Brain Atlas), at high levels in the hippocampus and cerebellum (Ruat et al., 1995). The neuronal function of CaSR however is largely unknown (Riccardi and Kemp, 2012). In cultured mouse hippocampal neurons and sympathetic neurons, blocking CaSR inhibits dendritic growth (Vizard et al., 2008). At the nerve terminal and cell bodies, CaSR activation suppresses a nonselective cation channel, inhibits evoked synaptic transmission, and activates spontaneous neurotransmitter releases (Chen et al., 2010; Phillips et al., 2008; Smith et al., 2004; Vyleta and Smith, 2011; Xiong et al., 1997).

Human CaSR mutations have been implicated in epilepsy ((Riccardi and Kemp, 2012) for a review). Hypothetically, the epilepsy could be because of alteration in serum [Ca^2+^] due to disruption in the PTH level, which leads to suppressing inhibitory neurons, activating excitatory neurons, or a more general effect on certain brain circuitry. Intriguingly, several reported CaSR mutations do not result in altered systemic PTH or serum Ca^2+^ level and yet are associated with seizure in affected individuals (Kapoor et al., 2008). How these mutations affect brain function is unknown. In this report, we found that the mutations disrupt CaSR’s ability to regulate the Na^+^-leak channel NALCN in neurons.

NALCN is an ~ 200 kDa protein highly conserved among animals. It shares sequence and structural similarities with the pore-forming α subunits of voltage-gated Ca^2+^ (Ca_v_) and Na^+^ (Na_v_) channels (Kang et al., 2020; Lee et al., 1999; Xie et al., 2020). The channel, as recorded in heterologous expression system and in neurons, is non-selective among cations and is largely voltage-independent (Eigenbrod et al., 2019; Funato et al., 2016; Hahn et al., 2020; Lee et al., 2019; Lu et al., 2007; Shi et al., 2016) (but see (Chua et al., 2020)). In mammalian brains, NALCN interacts with two large proteins UNC80 and UNC79, homologs of *C. elegans* proteins Unc-80 and Unc-79, respectively ((Ren, 2011) for a review). Affinity depletion of NALCN protein from brain lysates also depletes UNC79 and UNC80, suggesting the latter two are exclusively associated with NALCN and are auxiliary subunits of the NALCN complex (Wie et al., 2020). NALCN also associates with a smaller subunit NLF-1 (FAM155A) (Kschonsak et al., 2020; Xie et al., 2013). Mutations in humans and other animals in the complex result in developmental delay, hypotonia, epilepsy, lack of speech development, severe intellectual disability, breathing deficiency, disrupted circadian rhythm, altered sensitivity to anesthetics and premature death (Al-Sayed et al., 2013; Angius et al., 2018; Aoyagi et al., 2015; Bramswig et al., 2018; Campbell et al., 2018; Chong et al., 2015; Humphrey et al., 2007; Jospin et al., 2007; Koroglu et al., 2013; Kuptanon et al., 2019; Lear et al., 2013; Nash et al., 2002; Obeid et al., 2018; Perez et al., 2016; Pierce-Shimomura et al., 2008; Ren, 2011; Shamseldin et al., 2016; Stray-Pedersen et al., 2016; Takenouchi et al., 2018; Valkanas et al., 2016; Wie et al., 2020; Yeh et al., 2008).

In cultured hippocampal neurons and midbrain dopaminergic neurons, we and others have previously shown that lowering [Ca^2+^]_e_ increases NALCN-mediated Na^+^ leak channel current (I_NALCN_) and excites neurons (Lu et al., 2010; Philippart and Khaliq, 2018). This regulation of NALCN by Ca^2+^_e_ is G-protein-dependent and appears to involve CaSR, as CaSR agonists spermidine and neomycin suppresses the Ca^2+^ sensitivity of NALCN. NALCN is also regulated by other Gi/o-coupled receptors (Philippart and Khaliq, 2018). Whether CaSR-mediated NALCN regulation exists in other central nervous system (CNS) neurons and in peripheral neurons is unknown. In addition, the structural requirements on CaSR for the receptor’s coupling of Ca^2+^_e_ to the channel is not established. In this study, we used shRNA knockdown to demonstrate that CaSR is required for the regulation of NALCN by Ca^2+^_e_ in neurons. The ability to regulate NALCN by CaSR requires its Ca^2+^-sensing and, surprisingly, a domain in the C-terminus that is dispensable for the ability of the receptor to sense [Ca^2+^]_e_ and to couple it to other canonical targets. Finally, epilepsy-implicated mutations in this domain that do not lead to changes in the systemic [Ca^2+^] levels nevertheless disrupt the receptor’s ability to regulate NALCN, thus providing a novel mechanism by which the mutations cause epilepsy.

## RESULTS

### NALCN mediates Na^+^ leak in multiple brain regions and spinal cord

We have previously recorded the TTX-insensitive background Na^+^ leak currents (I_LNA_) from hippocampal neurons and found that I_LNA_ is largely mediated by NALCN, as knocking out NALCN in the mice eliminated most of the current (Lu et al., 2007). To test whether other CNS neurons also have NALCN-dependent I_LNA_, we measured I_LNA_ in cultured mouse cortical, ventral tegmental area (VTA) and spinal cord neurons. Because of its small sizes, we measured I_LNA_ by changes in the sizes of holding currents (at −68 mV) upon a drop in extracellular [Na^+^] from 140 mM to 14 mM (with 126 mM Na^+^ substituted by NMDG^+^, Figure 1A) (Lu et al., 2007). Similar to hippocampal neurons, all the types of CNS neurons cultured from wild-type (WT) mice have a non-inactivating background current I_LNA_ (Figure 1A, C). Such currents are largely absent in neurons cultured from the NALCN knockout (KO, NALCN^-/-^, Figure 1B, C). These data, together with those from others (Flourakis et al., 2015; Ford et al., 2018; Lu et al., 2007; Lutas et al., 2016; Philippart and Khaliq, 2018; Shi et al., 2016; Yeh et al., 2017), suggest that NALCN-mediated basal Na^+^ conductance is widely expressed in the CNS. The detection of I_LNA_ is consistent with the wide expression of the NALCN gene as demonstrated by in-situ RNA staining in the brain ((Lu et al., 2007) and Allen Brain Atlas).

**Figure 1.**
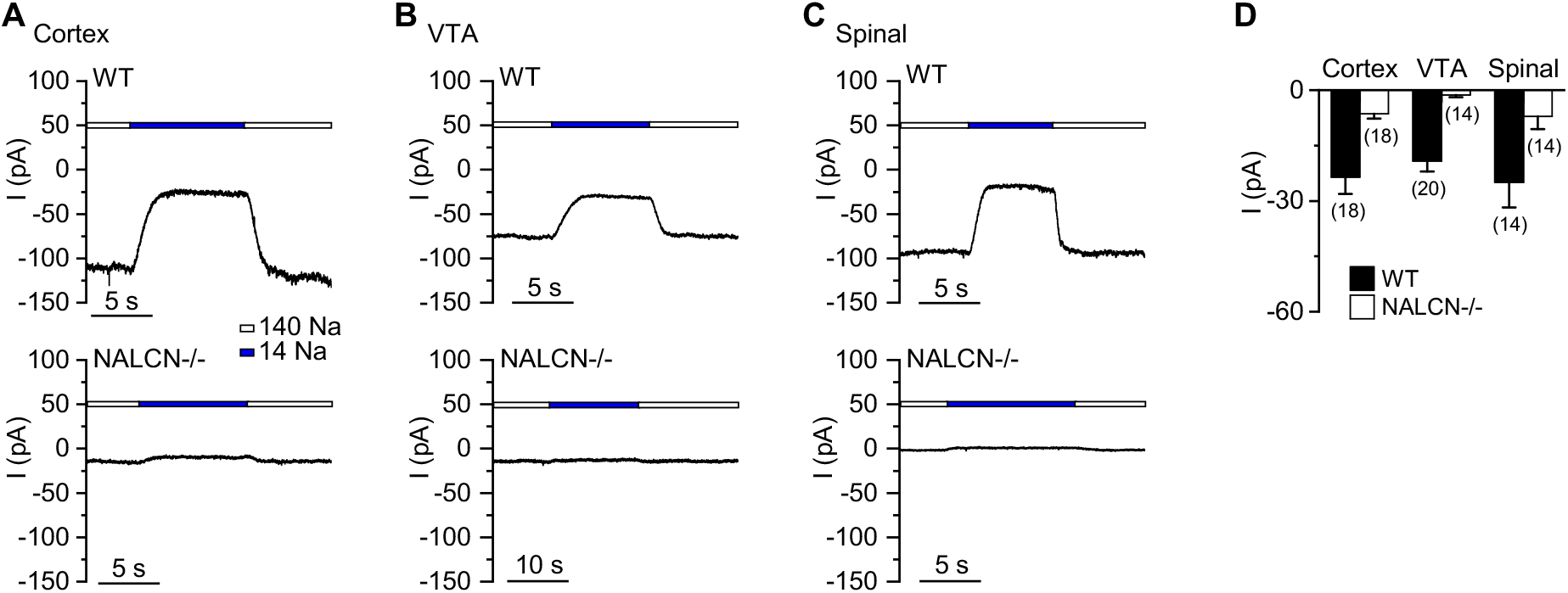
NALCN-dependent Na^+^-leak conductance is widely expressed the brain and spinal cord. Na^+^-leak currents (I_LNA_) were measured as changes in the sizes of holding currents (at −68 mV) upon a reduction of extracellular [Na^+^] from 140 mM to 14 mM. (**A-C**) Representative holding currents recorded from wild-type (WT, *upper* traces) and NALCN KO (NALCN-/-, *lower* traces) cortical (**A**), VTA (**B**) and spinal cord (**C**) neurons. (**D**) Average I_LNA_ sizes. Numbers of cells recorded are given in parentheses. Data is presented as mean ± S.E.

### Extracellular Ca^2+^ regulates NALCN-dependent Na^+^ currents in cortical, VTA and spinal cord neurons

In cultured hippocampal neurons, extracellular Ca^2+^ controls I_LNA_, and such control provides a way for Ca^2+^_e_ to regulate neuronal excitability (Lu et al., 2010). We tested whether Ca^2+^_e_ also regulates I_LNA_ in other CNS neurons by recording the Na^+^-dependent current before and after [Ca^2+^]_e_ was reduced from 2 mM to 0.1 mM (Figure 2A). Similar to that in hippocampal neurons, lowering [Ca^2+^]_e_ induced a Na^+^-dependent inward currents in wild-type neurons cultured from cortices (Figure 2A, D), VTA (Figure 2B, D) and spinal cord (Figure 2C, D). Strikingly, such low [Ca^2+^]_e_-induced current (I_LCA_) was essentially absent in the NALCN^-/-^ neurons (see also (Philippart and Khaliq, 2018) for dopaminergic neurons). These data suggest that both I_LNa_ and I_LCA_ are mediated by NALCN in the CNS neurons.

**Figure 2.**
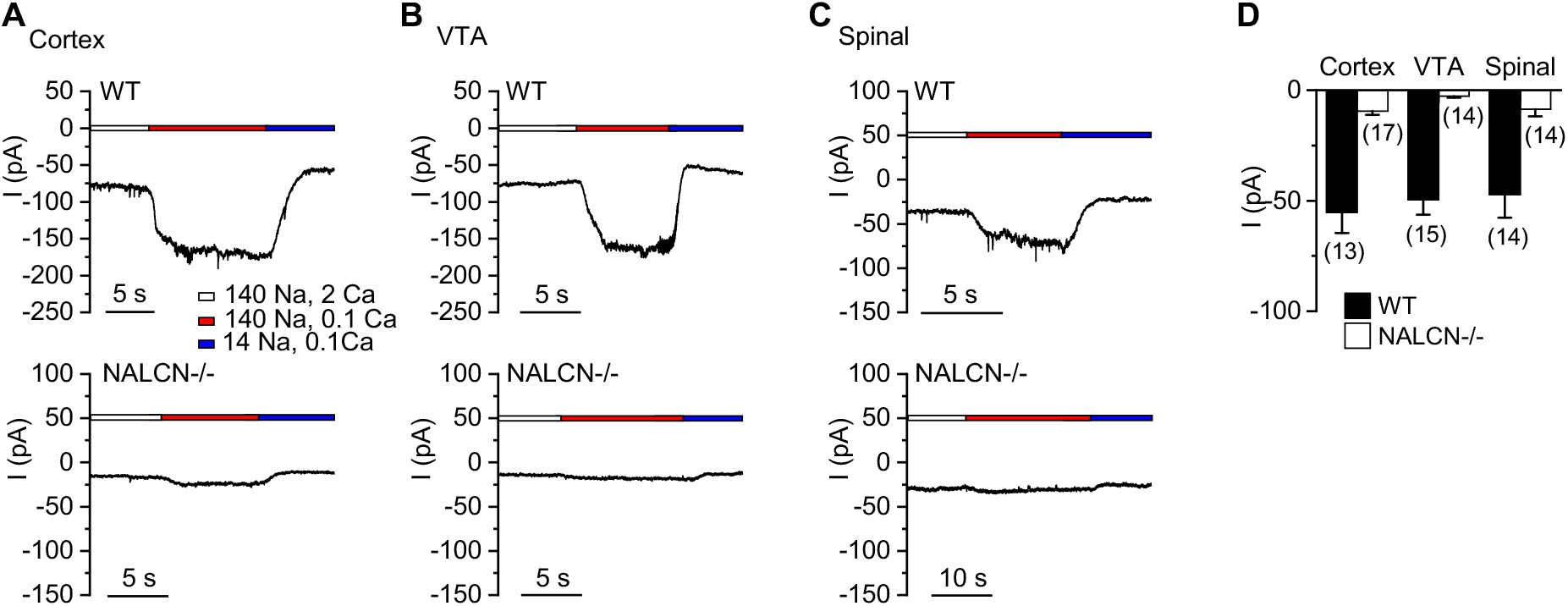
Extracellular Ca^2+^ regulates NALCN-dependent Na^+^ currents in the brain and spinal cord. Low [Ca^2+^]_e_-induced NALCN-dependent leak currents (I_LCA_) were measured as the changes in the sizes of holding currents (at −68 mV) when extracellular [Ca^2+^] was lowered from 2 mM (2 Ca^2+^) to 0.1 mM (0.1 Ca^2+^). In the end of each recording, bath was switch from the one containing 140 mM Na^+^ to one with 14 mM Na^+^ (126 mM Na^+^ replaced with NMDG) to ensure that the increase of holding currents in low [Ca^2+^] containing bath was not due to non-specific leak. (**A-C**) Representative currents recorded from a wild-type *(upper* traces) and *Nalcn* KO *(lower* traces) cortical (**A**), VTA (**B**) and spinal cord (**C**) neurons. (**D**) Averaged I_LCA_ sizes.

### Extracellular Ca^2+^ regulates NALCN-dependent Na^+^ currents in peripheral neurons

Like CNS neurons, peripheral neurons such as the dorsal root ganglion (DRG) neurons also have resting membrane potentials (RMPs) significantly depolarized to E_K_. Persistent Na^+^ conductance resulting from the window-conductance of voltage-activated Na^+^ channels Na_v_1.8 and Na_v_1.9 are present in the neurons (Herzog et al., 2001). However, DRG neurons in Na_v_1.8 and Na_v_1.9 knockout mice are not hyperpolarized compared to the WT (Priest et al., 2005), suggesting the presence of other subthreshold Na^+^ conductance. To test whether NALCN contributes to the conductance, we compared I_LNa_ between WT and NALCN^-/-^ DRG neurons. WT DRG neurons had large TTX-resistant I_LNA_ (Figure 3A, C). Such current was absent in the NALCN^-/-^ neurons. In WT, the current was also potentiated by lowering [Ca^2+^]_e_, but such low [Ca^2+^]_e_ -mediated current was largely absent in the KO (Figure 3B, D). Together, our data suggest that in both CNS and PNS neurons, NALCN contributes the basal Na^+^ conductance and it is regulated by Ca^2+^_e_.

**Figure 3.**
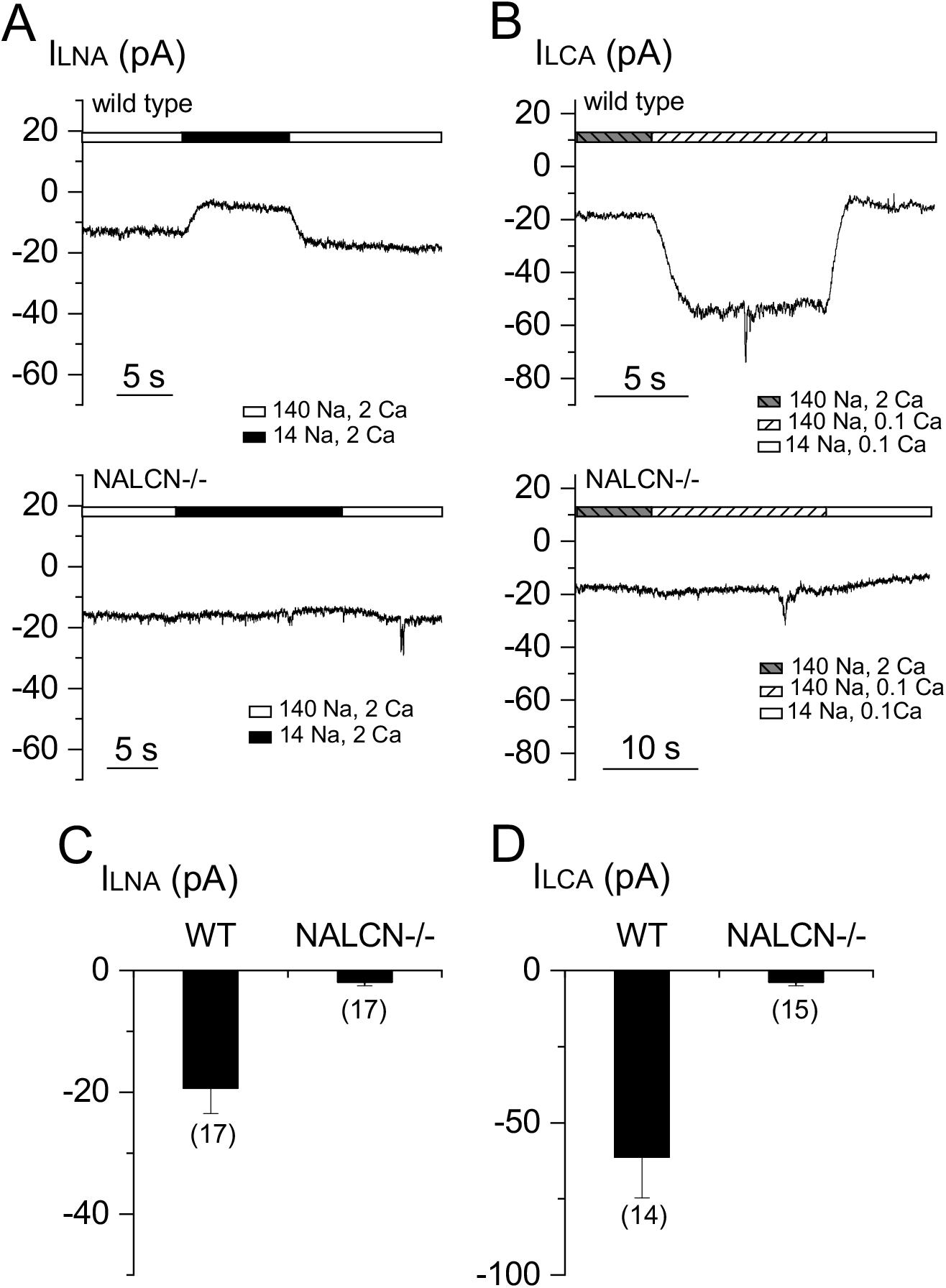
Extracellular Ca^2+^ regulates Na^+^-leak currents in peripheral neurons. (**A, B**) Representative I_LNA_ (recorded with 140 mM Na^+^-containing bath) (**A**) and I_LCA_ (**B**) recorded in DRG neurons cultured from wild type (WT, *upper panels*) and NALCN knockout (NALCN-/-, *lower panels*) mice. (**C, D**) Averaged sizes of the I_LNA_ (**C**) and I_LCA_ (**D**).

### CaSR is required for the Ca^2+^ sensitivity of NALCN

How NALCN is sensitive to Ca^2+^_e_ is not well understood. Ca^2+^_e_ inhibits cation channels such as Ca_v_s and TRPs by blocking the channel pore (Owsianik et al., 2006; Yang et al., 1993). The Ca^2+^ sensitivity of neuronal NALCN under quasi-physiological conditions does not appear to involve such a pore-blocking mechanism (Lu et al., 2010). In hippocampal neurons, CaSR agonists spermidine and neomycin inhibit I_LCA_, implicating CaSR in the regulation of NALCN (Lu et al., 2010; Xiong et al., 1997). To further test whether CaSR gene expression is required for NALCN’s Ca^2+^ sensitivity in neurons, we designed an shRNA specifically against the mouse CaSR (mCaSR) gene (Figure 4A). In a heterologous expression system, the shRNA knocked down ~90% of CaSR protein (Figure 4A, B). Transfection of the shRNA, but not a control with sequence mismatches, into cultured hippocampal neurons, eliminated I_LCA_ (Figure 4C, D). These data suggest that CaSR gene expression is required for the Ca^2+^ sensitivity of NALCN in neurons.

**Figure 4.**
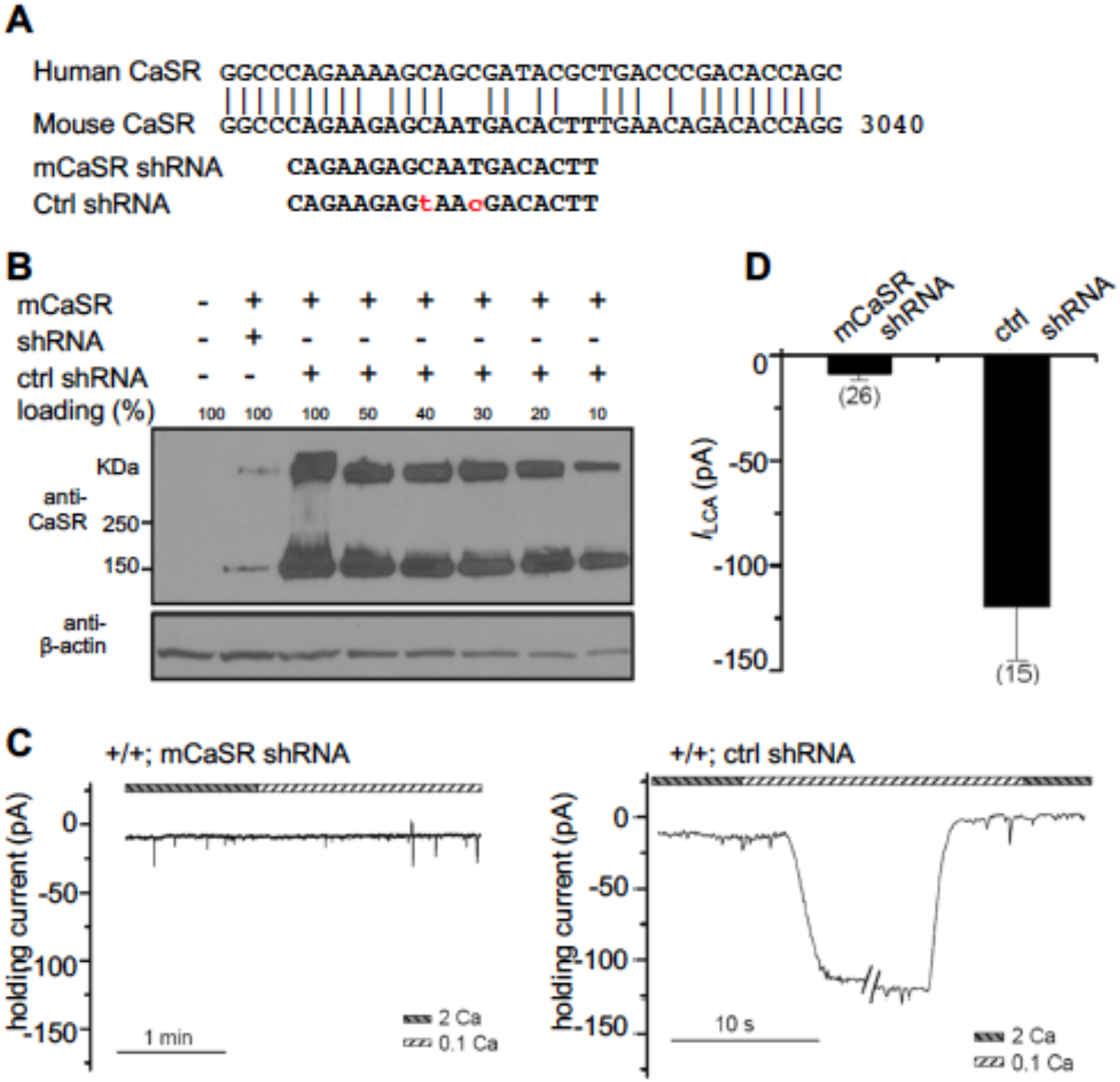
CaSR is required for the regulation of NALCN currents by extracellular Ca^2+^ in neurons. (**A**) Design of an shRNA specific to the mouse CaSR (mCaSR). *Upper:* alignment of human (h) and mouse (m) CaSR sequences in the region targed by the shRNA. *Lower:* sequence of the shRNA target against mCaSR and that of a control shRNA with mismatches at two base pairs. (**B**) Western-blots showing the knockdown efficiency of the shRNA. CaSR protein levels were detected with cell lysates from HEK293T cells co-transfected with mouse CaSR cDNA (mCaSR) and the shRNA against mCaSR or the missense shRNA as control. Decreasing amounts of total cell lysates from the “mCaSR + missense shRNA”-transfected cells were loaded to determine the knockdown efficiency. “100%” indicates the amount equal to that used from the non-transfected cells. Anti-β-actin was used as the loading control. The upper bands in the blot with anti-CaSR represent protein dimmers. (**C**) Representative I_LCA_ recordings in WT hippocampal neurons transfected with shRNA against mouse CaSR *(left* panel) or the missense shRNA as control *(right* panel). (**D**) Summary of the I_LCA_ current sizes.

### CaSR’s Ca^2+^ -sensing ability is required for its regulation of NALCN

To define the structural requirements of CaSR in the regulation of NALCN in neurons, we developed a “knockdown-and-reconstitution” system where wild-type or mutant human CaSR (hCaSR) was transfected into mouse hippocampal neurons in which the endogenous mCaSR was depleted with a co-transfected shRNA specifically against mCaSR. HCaSR is highly similar to mCaSR at the amino acid level (93% identity and 96% similarity), but differs from mCaSR at the nucleotide level. The target sequences of the shRNA designed against mCaSR do not match those of hCaSR (with 6 mismatches, Figure 4A). As such, the hCaSR used for transfection is resistant to the mCaSR-specific shRNA. Transfecting wild-type hCaSR completely restored I_LCA_ in the mCaSR-knocked down neurons (Figure 4C, Figure 5A, C).

**Figure 5.**
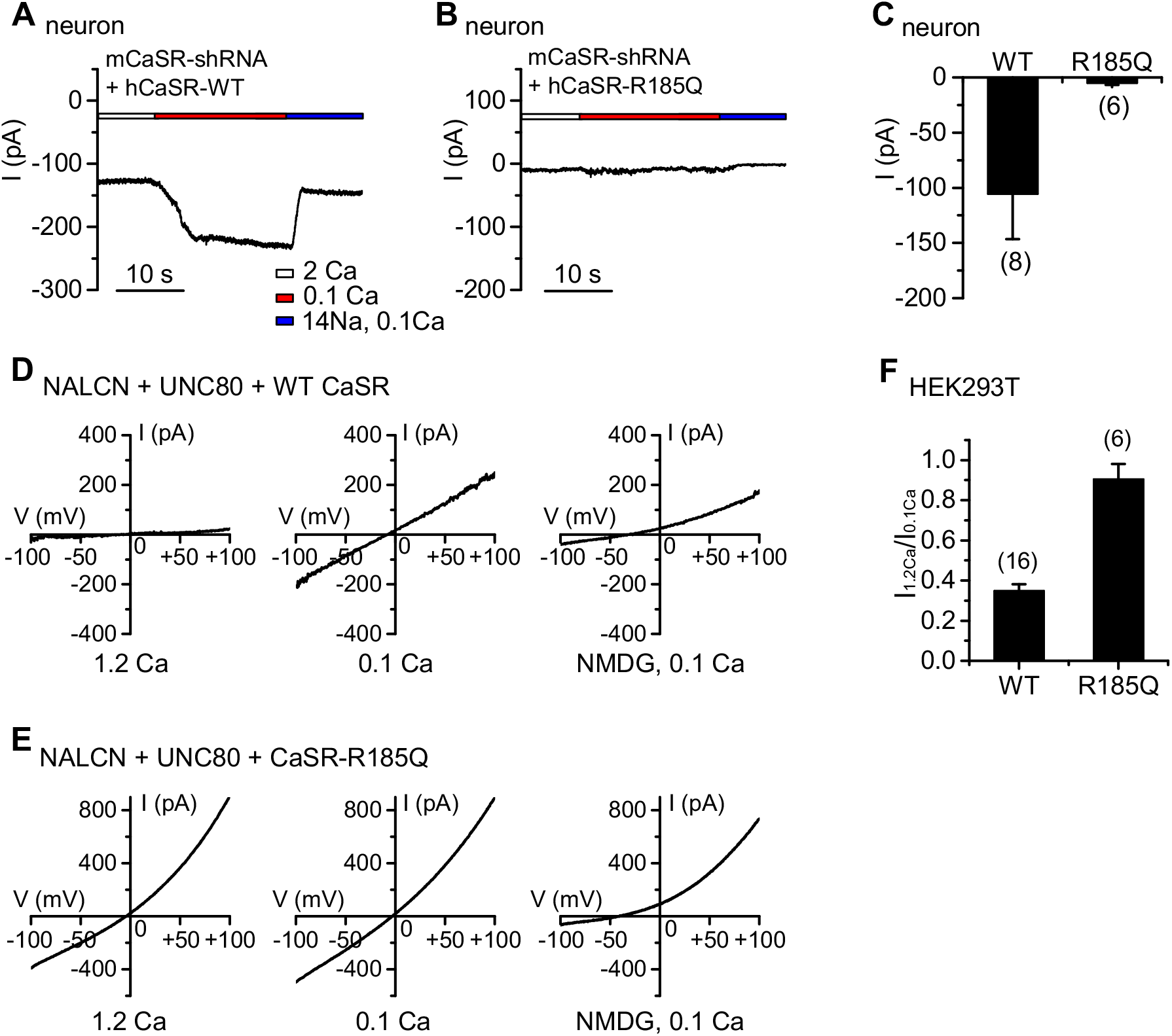
CaSR’s Ca^2+^-sensing ability is required for the receptor’s regulation of NALCN currents. (**A-C**) Representative I_LCA_ recordings in WT mouse hippocampal neurons transfected with a wild-type (**A**) or R185Q mutant (**B**) human CaSR cDNA. The endogenous mCaSR was knocked down with co-transfection of the mCaSR-specific shRNA. Summary of I_LCA_ sizes are in (**C**). (**D-F**) I_NALCN_ recorded from HEK293T cells co-transfected with NALCN, UNC80 and a wild-type human CaSR (**D**) or the R185Q mutant (**E**), recorded with 1.2 mM (1.2 Ca, *left panels)* or 0.1 mM (0.1 Ca, *middle panels)* Ca^2+^ in the bath. In the end of recording, bath was perfused with one in which Na^+^ and K^+^ were replaced with NMDG *(right panels).* A ramp protocol was used (−100 mV to +100 mV in 1 s, V_h_ = 0 mV). In (**F**), the Ca^2+^-sensitivity of I_NALCN_ is represented as the ratio of the sizes of current (at −100 mV) recorded with bath containing 1.2 mM (I_1.2Ca_) and that recorded with 0.1 mM Ca^2+^-containing bath (I_0.1Ca_).

CaSR detects changes in [Ca^2+^]_e_ through conformational changes upon ligand-binding to its extracellular ligand binding domain, and transduces the signal to intracellular targets such as phospholipase C (PLC) and protein kinases ((Brown et al., 1993; Hofer and Brown, 2003) for reviews). An hCaSR bearing a hyperparathyroidism- and hypercalcemia-associated point mutation R185Q in the receptor’s extracellular domain is unable to sense [Ca^2+^]_e_ (Bai et al., 1997). In contrast to the WT hCaSR, the R185Q mutant was unable to restore [Ca^2+^]_e_-sensitive I_NALCN_ in neurons in which the endogenous mCaSR was knocked down (Figure 5B, C).

NALCN’s Ca^2+^_e_-sensitivity can also be reconstituted in HEK293T cells by co-transfecting CaSR, Src529 (a constitutively active Src mutant) and UNC80 (Lu et al., 2009; Stray-Pedersen et al., 2016). In contrast to WT, the R185Q mutant was unable to reconstitute NALCN’s Ca^2+^ sensitivity (Figure 5D-F). These data suggest that CaSR’s ability to sense Ca^2+^_e_ is required for the coupling of changes in [Ca^2+^]_o_ to NALCN by the receptor.

### CaSR’s C-terminus, which is dispensable for Ca^2+^-sensing, is required for the NALCN regulation

We further tested whether the ability of CaSR to couple Ca^2+^_e_ to its known intracellular targets is sufficient to couple Ca^2+^_e_ to NALCN. CaSR contains a large intracellular C-terminus, 216 residues in the human isoform. The C-terminus is highly conserved among mammals (Figure 6A), but its function is not well established. Compared to the wild-type (WT), hCaSR mutants lacking as large as 190 amino acids in the C-terminus (deletion after T888) have apparently equal or better ability for surface expression, receptor dimerization, Ca^2+^ sensing, stimulation of IP3 production and activation of the MAPK pathway (Bai et al., 1998; Chang et al., 2001; Gama and Breitwieser, 1998; Ray et al., 1997; Zhang and Breitwieser, 2005; Zhuang et al., 2012), which are some of the most well characterized cellular functions of CaSR.

**Figure 6.**
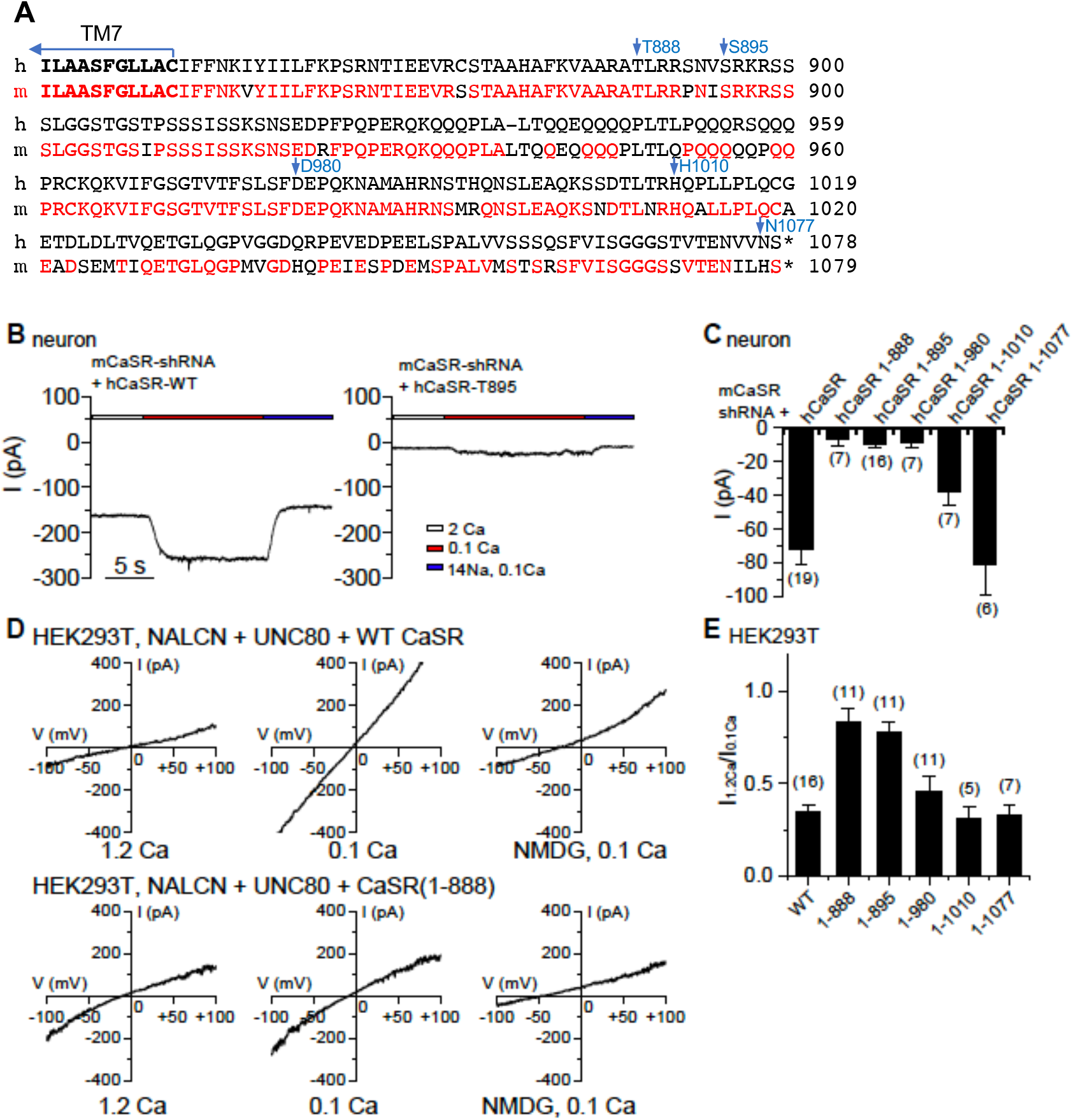
The C-terminus of CaSR is required for its ability to regulate NALCN. (**A**) Sequence alignment of the C-termini of human (h) and mouse (m) CaSR. Mouse sequences conserved with those in human are in red. (**B**) Representative I_LCA_ recorded in mouse hippocampal neurons co-transfected with mCaSR shRNA and full-length hCaSR or a mutant truncated after T895 (hCaSR-T895). (**C**) Averaged amplitudes of I_LCA_ recorded in mouse hippocampal neurons co-transfected with mCaSR shRNA and full-length hCaSR or mutant hCaSR with truncation as indicated. (**D, E**) I_NALCN_ recorded from HEK293T cells co-transfected with NALCN, UNC80 and a wild-type human CaSR or a mutant truncated at T888, recorded with 1.2 mM (1.2 Ca, *left panels* in **D**) or 0.1 mM (0.1 Ca, *middle panels)* Ca^2+^ in the bath. In the end of recording, bath was perfused with one in which Na^+^ and K^+^ replaced with NMDG *(right panels)* A ramp protocol was used (−100 mV to +100 mV in 1 s, V_h_ = 0 mV). (**E**) Ca^2+^-sensitivity of I_NALCN_ is plotted as the ratio of the sizes of current (at −100 mV) recorded with bath containing 1.2 mM (I_1.2Ca_) and that recorded with 0.1 mM Ca^2+^-containing bath (I_0.1Ca_). The ratio for WT was replotted from Figure 5F for comparison.

Remarkably, hCaSR truncated at T888 (CaSR1-888) almost completely lacked the ability to confer NALCN’s Ca^2+^ sensitivity in neurons (Figure 6C). Additional truncational mutants suggest that CaSR retains its ability to regulate NALCN in neurons when containing amino acids 1-1010 or 1-1077, but the ability is significantly reduced when CaSR is truncated to amino acid 895 or 980 (Figure 6B, C).

We further tested the importance of CaSR’s C-terminus in heterologous expression system. Similar to those tested in neurons using the knockdown-and-reconstitution system, the Ca^2+^-sensitivity of NALCN currents (represented as the ratio of current amplitudes measured under 1.2 and 0.1 mM [Ca^2+^]_e_, I_1.2Ca_/I_0.1Ca_) remained apparently intact in cells co-expressing hCaSR with the C-terminus deleted up to amino acid 1010, but was largely diminished when hCaSR was further truncated to amino acid 980 (Figure 6D, E). These results suggest that the C-terminal region of aa 888-1010 plays an essential role in regulating NALCN. CaSR without this region, though able to sense Ca^2+^_e_ and to couple it to other known signaling cascades such as the generation of IP3 and the activation of MAPK, is unable to couple Ca^2+^_e_ to NALCN,

### Epilepsy-implicated CaSR mutations disrupt the receptor’s ability to regulate NALCN

Given the differential requirements of CaSR’s C-terminus in the receptor’s ability to couple Ca^2+^_e_ to NALCN and that to the other targets such as PLC and MAPK, we searched human CaSR mutations that might specifically affect the receptor’s coupling to a subset of targets. Many human CaSR mutations are implicated in epilepsy, but the underlying mechanisms are not well understood (Pidasheva et al., 2004). We tested a panel of CaSR mutations (E354A, R898Q and A988V) found in epileptic patients who do not have abnormal systemic Ca^2+^ or PTH levels (Kapoor et al., 2008). The apparent normal levels of serum [Ca^2+^] indicate normal functional coupling by the CaSR from Ca^2+^_e_ to intracellular targets such as PLC in the generation of PTHs to maintain the systemic [Ca^2+^] level. Each of the mutations, however, reduced the inhibitory effect of CaSR on I_NALCN_ when reconstituted in HEK293T cells (Figure 7A-C) (I_1.2Ca_/I_0.1Ca_: A988V, 0.70 ± 0.04, n = 10; R898Q, 0.64 ± 0.05, n = 10; E354A, 0.79 ± 0.04, n = 9), even though the protein expression levels of the mutants were comparable with that of the wild-type (Figure 7D).

**Figure 7.**
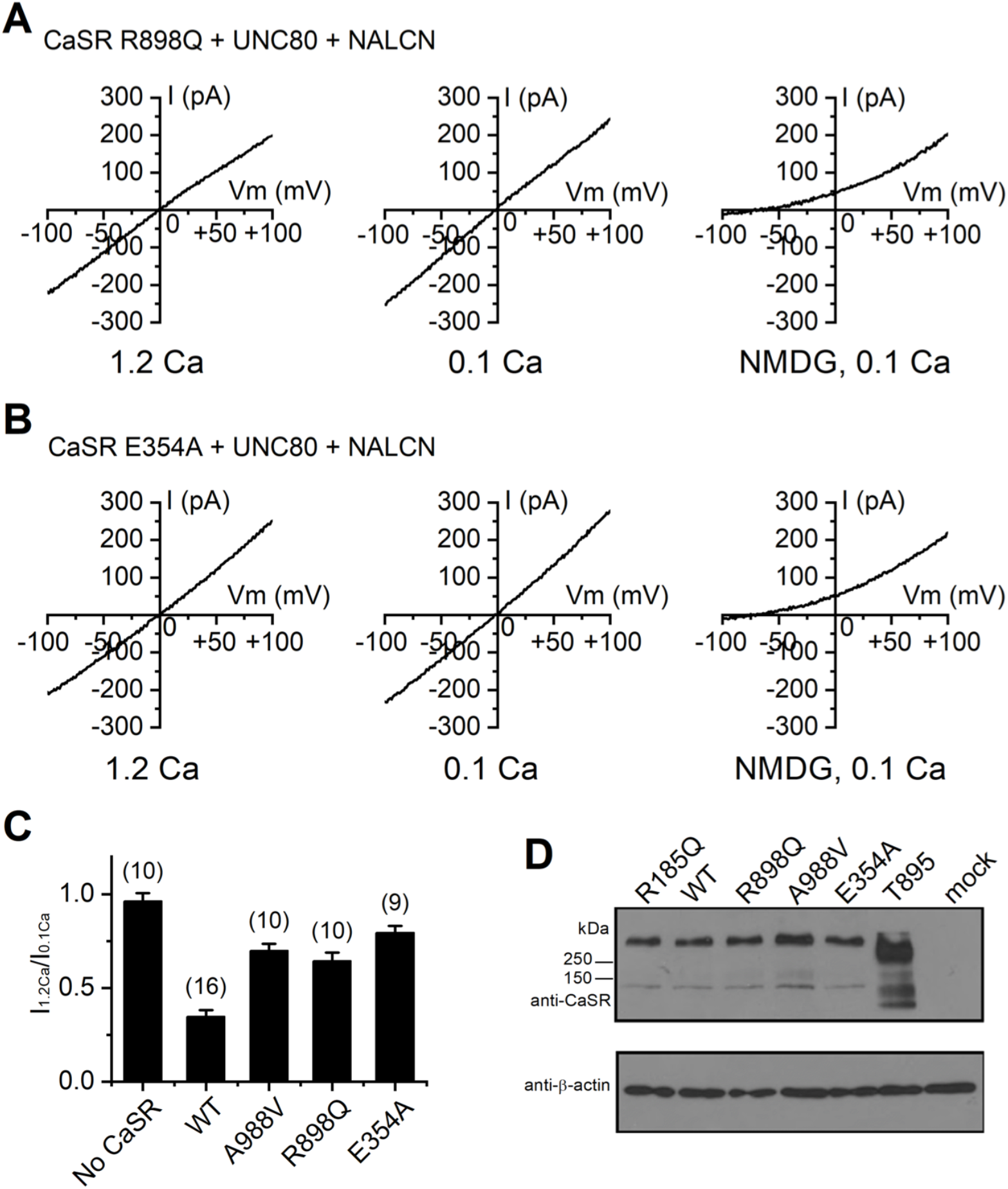
Epilepsy-associated mutations in CaSR disrupt CaSR’s ability to regulate NALCN in HEK293T cells. (**A, B**) Representative I_NALCN_ recorded from HEK293T cells co-transfected with NALCN, UNC80 and human CaSR mutants R898Q (**A**) or E354A (**B**), recorded with 1.2 mM (1.2 Ca, *left panels)* or 0.1 mM (0.1 Ca, *middle panels)* Ca^2+^ in the bath. In the end of recording, bath was perfused with one in which Na^+^ and K^+^ replaced with NMDG *(right panels*). A ramp protocol was used (−100 mV to +100 mV in 1 s, V_h_ = 0 mV). (**C**) Averaged Ca^2+^-sensitivity of I_NALCN_ reconstituted with WT or mutant CaSR, as represented as the ratio of the sizes of current (at −100 mV) recorded with bath containing 1.2 mM (I_1.2Ca_) and that recorded with 0.1 mM Ca^2+^-containing bath (I_0.1Ca_). The ratio for WT was replotted from Figure 5F for comparison. (**D**) Western blot with anti-CaSR or anti-β-actin (for loading control) of lysates of HEK293T cells transfected with empty vector (mock), WT or mutant hCaSR as indicated. The carboxy-terminal truncated CaSR mutant (CaSR 1-895) gave rise to apparently higher level of protein when the same amount of DNA (2 μg) as the wild-type was used for transfection. Recordings were also done with 1/3 amount of CaSR 1-895 (0.7 μg) as normally used (2 μg); still, no obvious suppression of the NALCN current was observed (I1.2Ca/I0.1Ca = 95.9 + 17.6 %; n = 8).

We also tested the epilepsy-associated CaSR mutations in hippocampal neurons. In cultured mouse hippocampal neurons in which the native mCaSR was knocked down, transfection of the mutant hCaSRs only partially restored I_NALCN_’s Ca^2+^ sensitivity, in contrast to those transfected with WT hCaSR (Figure 8).

**Figure 8.**
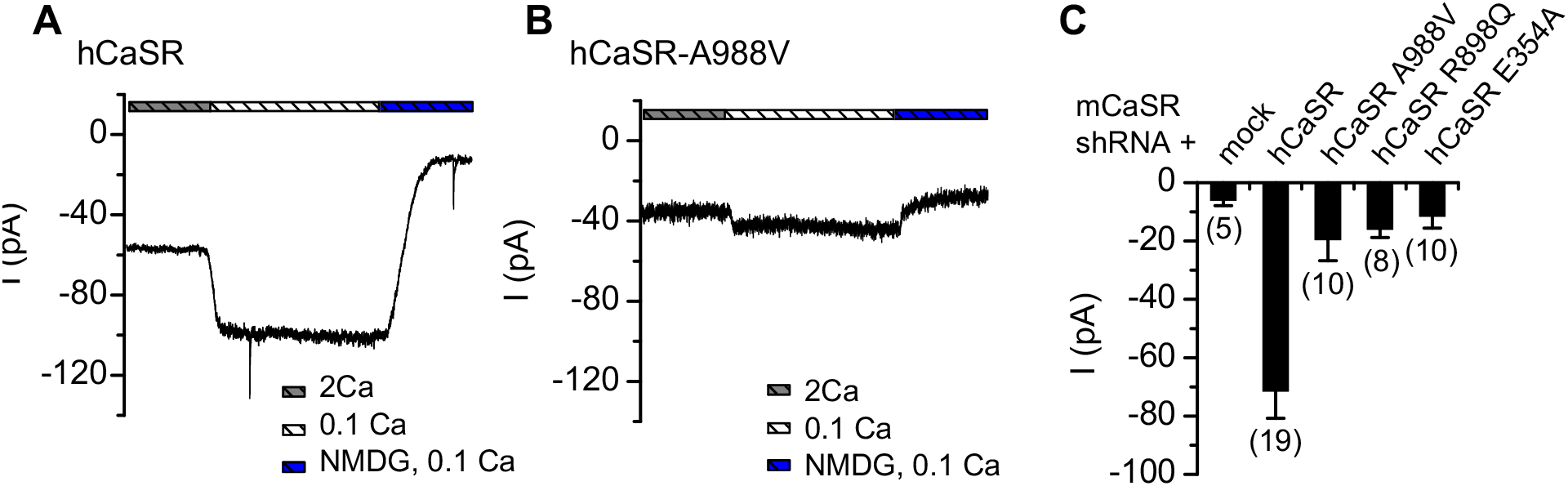
Epilepsy-associated mutations in CaSR’s C-terminus disrupt I_LCA_ in neurons. (**A, B**) Representative I_LCA_ recordings in WT hippocampal neurons transfected with the wild-type (**A**) or A99V mutant (**B**) human CaSR cDNA. The endogenous mouse CaSR was knocked down with co-transfection of the mouse CaSR-specific shRNA. Summary of the I_LCA_ current sizes with transfection of wild-type and the epilepsy-associated mutant hCaSR as indicated are in (**C**).

## DISCUSSION

We have shown that the NALCN-dependent basal Na^+^ conductance is widely present in the central and peripheral nervous systems. Extracellular Ca^2+^ also regulates the conductance via a CaSR-dependent mechanism in neurons. In addition to the Ca^2+^-sensing ability of CaSR, a C-terminal domain of CaSR located between amino acids 888-1010, while not essential for the canonical functions of the receptor, is required for the receptor’s coupling of Ca^2+^_e_ to NALCN. The importance of such coupling is demonstrated by the epilepsy-implicated CaSR mutations that specifically disrupt the receptor’s ability to regulate NALCN while maintaining the receptor’s functional coupling to other conventional targets in the maintenance of systemic Ca^2+^ and PTH levels.

CaSR is coupled to Gqa and its activation leads to increases in intracellular IP3 and Ca^2+^ levels (Hofer and Brown, 2003). This well-established function of CaSR appears to be insufficient for the receptor’s ability to regulates NALCN current. The 183 amino acids of the distal carboxy terminus of CaSR are well conserved among vertebrates, suggesting that they have physiological function, but they are not required for the receptor’s ability to sense [Ca^2+^]_e_ changes and to trigger IP3 production (Chang et al., 2001; Gama and Breitwieser, 1998; Ray et al., 1997). In contrast, this segment is essential for the ability of CaSR to suppress I_NALCN_ (Figure 6), suggesting a novel function for this intracellular tail. Consistent with this proposed function of the carboxy terminus of CaSR in NALCN regulation, but not in the [Ca^2+^]_e_ detection in the parathyroid gland, the R898Q mutation in the region found in human patients does not lead to abnormality in whole-body Ca^2+^ homeostasis; it does, however, lead to a deficiency in I_LCA_ and is associated with heritable epilepsy (Kapoor et al., 2008). Future studies will need to investigate the structural basis underlying this domain’s involvement in the coupling between CaSR and the NALCN channel complex.

## EXPERIMENTAL PROCEDURES

### Animals

Animal uses were approved by the University of Pennsylvania IACUC. The development of NALCN knockout was previously described (Lu et al., 2007). P0 pups were obtained from matings between heterozygous. Littermates were used as controls.

### DNA constructs

The cDNA constructs encoding NALCN (rat), UNC80 (mouse, in vector pcDNA3.1(+)) and Src were previously described (Lu et al., 2007; Lu et al., 2009; Lu et al., 2010). The NALCN construct was made in a vector based on pTracer-CMV2 (Invitrogen) and modified to also express eGFP under a separate promoter. Human CaSR was amplified from an IMAGE EST clone (ID#8327704) and subcloned into the NotI and XbaI sites of a vector based on pTracer-CMV2 modified to also express mCherry RFP under a separate promoter for the identification of transfected cells. CaSR mutants were constructed by introducing point mutations using PCR methods as follows: R185Q (nt554/G mutated to A; with the first nucleotide (nt) in the open reading frame numbered as 1), E354A (nt1065/A to C), R898Q (nt 2693/G to A), A988V (nt2963/C to T), CaSR1-895 (nt2686-2688/CGC to TGA; amino acid 896 mutated to a stop codon). The mouse CaSR open reading frame used for the test of shRNA efficiency was amplified from mouse kidney cDNA and subcloned into the HindIII and SacII sites of peGFP-N1. The shRNA constructs were cloned into the GFP-containing vector pG-Super (Kojima et al., 2004) (a gift from Dr. Tatyana Svitkina) cut with BglII and HindIII, according to the manual of pSuper vector, with targeting sequence of CAGAAGAGCAATGACACTT. The mis-match control shRNA has a target sequence of CAGAAGAGtAAcGACACTT, with two mismatches to the mCaSR sequence. All DNA constructs were confirmed with sequencing.

### Cell culture and transfection

All the cells were maintained in humidified incubators at 37°C and 5% CO2. HEK293T cells (from ATCC) were cultured in DMEM (Gibco) medium supplemented with 1x penicillin-streptomycin (Pen-Strep, Gibco) and 10% FBS (Atlanta biologicals and BioWhittaker). Mouse hippocampal and spinal cord neurons were cultured from P0 as previously described (Lu et al., 2007; Lu et al., 2010). Briefly, dissected tissues were digested with papain (Worthington) and plated on poly-L-lysine-coated glass coverslips in 35-mm dishes at a density of ~3–4 ×10^5^ cells/dish, in 80% DMEM (Lonza), 10% Ham’s F-12 (Lonza), 10% bovine calf serum (iron supplemented, Hyclone) and 0.5x Pen-Strep. Medium was changed the next day to Neurobasal A medium supplemented with 2% B-27, 0.5x Pen-Strep, and 1x Glutamax (all from Gibco). VTA neurons were cultured similar to hippocampal neurons except that neurons were plated in the Neurobasal A-based medium on the first day. DRGs were dissected from P10 pups in ice-cold DMEM (Gibco) and were digested for 60 min at 37°C in a 35-mm dish with 1.5 ml DMEM, 4 mg collagenase type 2 (Worthington) and 1.5 mg trypsin (Worthington). Digestion was stopped with 1 ml culture medium containing 80% DMEM (Lonza), 10% BCS (Hylcone), 10% Ham’s F12 (Lonza), 0.5% Pen-Strep (Gibco). Cells were dissociated by pipetting up and down, and were plated onto poly-lysine coated coverslips in culture medium. Medium was changed the next day and once a week after. Cytosine arabinoside (ARAC, Sigma) was added at 5 μM to the medium when needed to suppress the growth of fibroblast.

DNA transfections were performed using Lipofectamine 2000 (Invitrogen) according to the manufacture’s instruction. Recordings were done 48-72 hr later. For I_LCA_ recording in HEK293T cells, cells (~90% confluency) plated on a 35-mm dish were transfected with 0.5 μg NALCN, 0.5 μg UNC80, 0.5 μg Src529 (a constitutively activated Src kinase), and 2 μg CaSR cDNA unless otherwise stated. In control experiments where one or more constructs were not included, an equal amount of empty vector DNA was added to ensure that all the transfections had the same amount of DNA. Cells were re-plated on the day of recording. Only cells with moderate level of both GFP and RFP fluorescence signals were selected for recordings. Neurons between DIV 5 and 7 were used for transfection. For shRNA knockdown in hippocampal neurons, 3 μg shRNA DNA was used for a 35 mm dish. For knockdown-and-reconstitution in neurons, 2 μg shRNA DNA and 1 μg CaSR DNA was co-transfected. DNAs used for neuronal transfection were endotoxin-free.

### Western blot

HEK293T cells in 35 mm dishes were lysed by incubation at 4°C for 1 hr in 300 μl RIPA buffer (50 mM Tris-HCl, 150 mM NaCl, 1% NP-40, 0.5% (w/v) deoxycholate, 0.1% (w/v) SDS, pH 7.4) supplemented with a protease inhibitor cocktail (PIC, Roche). After centrifuging for 30 min at 20,000 g, the supernatant was transferred to a fresh tube.

Protein electrophoresis was performed with 10% SDS-PAGE gel in Tris-glycine-SDS running buffer or 4-12 % Bis-Tris gradient gels in MOPS-SDS running buffer (Invitrogen). Proteins were transferred onto polyvinylidene difluoride (PVDF) membranes. After being blocked with 5% nonfat dry milk in PBS with 0.1% Tween-20 (PBST), membranes were incubated with primary antibodies (1 μg/ml) at 4°C overnight. Following incubation with horseradish-peroxidase-labeled secondary antibodies for 1 hr at room temperature, membranes were developed with SuperSignal West Pico ECL or SuperSignal West Dura ECL. The anti-CaSR antibody (monoclonal, against aa 15-29) was from Santa Cruz (cat #Sc-47741).

### Electrophysiology

For HEK293T cell recordings, the pipette solution contained (in mM) 150 Cs, 120 Mes, 10 NaCl, 10 EGTA, 4 CaCl_2_, 0.3 Na_2_GTP, 2 Mg-ATP, 10 HEPES (pH 7.4, osmolarity of ~300 mOsm/L). Bath contained (in mM) 150 NaCl, 3.5 KCl, 1 MgCl_2_, 10 HEPES, 20 glucose, and CaCl_2_ as indicated (pH 7.4 with 5 NaOH, ~320 mOsm/L). In the NMDG-containing bath, Na^+^ and K^+^ were replaced with NMDG^+^. The inclusion of ~ mM Mg^2+^ in both pipette and bath solutions minimizes the contamination of the endogenous Mg^2+^-inhibited TRPM7 currents present in HEK293 cells (Chokshi et al., 2012; Chubanov et al., 2012).

For recording from neurons, pipette solution contained (in mM) 120 CsCl, 4 EGTA, 2 CaCl_2_, 2 MgCl_2_, 4 Mg-ATP, 0.3 Tris-GTP, 14 phosphocreatine (di-tris salt) and 10 HEPES (pH 7.4). The 140 mM Na^+^-containing bath contained (in mM) 140 NaCl, 5 KCl, 2 (or 0.1) CaCl_2_, 1 MgCl_2_, 6 glucose, 2 CsCl and 10 HEPES (pH 7.4). In the 14 mM Na-containing bath, 126 mM NaCl was replaced with 126 mM Tris-Cl. To block voltage-gated Na^+^ channels and synaptic currents, TTX (1 μM), APV (10 μM), bicuculline (20 μM) and CNQX (20 μM) were added to the bath.

Patch clamp recordings were performed using an Axopatch-200A and a MultiClamp 700B amplifier controlled with Clampex 9.2 or Clampex 10 software (Axon). Signals were digitized at 2–10 kHz with a Digidata 1322A or 1440 digitizer. Liquid junction potentials (estimated using Clampex software) were corrected offline.

## ACKNOWLEDGMENTS

We thank members of the Ren lab for help and discussions. The work was supported, in part, by NIH grants 1R01NS074257 and 1R01NS055293 (to D.R.).

